# Parallel Capsule Net for Ischemic Stroke Segmentation

**DOI:** 10.1101/661132

**Authors:** MD Sharique, Bondi Uday Pundarikaksha, Pradeeba Sridar, R S Rama Krishnan, Ramarathnam Krishnakumar

## Abstract

Stroke is one of the leading causes of disability. Segmentation of ischemic stroke could help in planning an optimal treatment. Currently, radiologists use manual segmentation, which can often be time-consuming, laborious and error-prone. Automatic segmentation of ischemic stroke in MRI brain images is a challenging problem due to its small size, multiple occurrences and the need to use multiple image modalities. In this paper, we propose a new architecture for image segmentation, called Parallel Capsule Net, which uses max pooling in every parallel pathways along with dense connections between the parallel layers. We hypothesise that the spatial information lost due to max pooling in these layers can be retrieved by the use of such dense connections. In order to combine the information encoded by the parallel layers, outputs of the layers are concatenated before upsampling. We also propose the use of a modified loss function which consists of a regional term (Generalized Dice loss + Focal Loss) and a boundary term (Boundary loss) to address the problem of class imbalance which is prevalent in medical images. We achieved a competitive Dice score of 0.754, on ISLES SISS data set, compared to a score of 0.67 reported in earlier studies. We also obtained a Dice score of 0.902 with another popular data set, ATLAS. The proposed parallel capsule net can be extended to other similar medical image segmentation problems.

## 1 Introduction

The ischemic stroke which accounts for 87% of stroke cases is the most common cerebrovascular disease [1]. Ischemic stroke is caused by the obstruction of cerebral blood supply resulting in tissue hypoxia which progresses through several stages such as acute, subacute and chronic [2]. The lesions are qualitatively assessed by the use of multiple magnetic resonance imaging sequences (DWI, FLAIR, T2) as part of the clinical workflow. Currently, in the context of treatment decision and stroke research, the lesion is manually segmented by the radiologists that is often time consuming and error-prone.

There are a few challenges which are currently faced due to automatic segmentation of stroke lesions. Firstly, the lesion appearance changes considerably over time within a given sub stage of disease progression. Secondly, the lesion can be very small and can have multiple occurrences in the brain. Thirdly, several other common pathologies like small vessel ischemic disease(prevalent in older hypertensive and diabetic patients) and multiple sclerosis can have similar morphologic appearance as ischemic stroke on MR [2].

The Ischemic Stroke Lesion Segmentation (ISLES) 2015 challenge [2] had two sub-challenges: Sub-Acute Stroke Lesion Segmentation Challenge (SISS) and Stroke Perfusion Estimation (SPES). DeepMedic [3] (*Konstantinos Kamnitsas et al.*) used multi-scale 3D CNN with fully connected CRFs achieving the highest Dice score of 0.59 in SISS. *Zhiyang Liu et al.* [4] proposed a residual-structured fully convolutional network (Res FCN) that does 2D slice-based segmentation using dice coefficient as the loss function. *Chen et al.* [5] made use of EDD Net (an ensemble of DeconvNets) and MUSCLE Net (Multiscale Convolutional Label Evaluation Net) to segment and refine the lesion by removing False positives. By embedding a residual unit into a U shaped network, *Liangliang Liu et al*. [6] achieved an average dice score of 88.43 in SPES. Based on DenseNets, [7] proposed an architecture that uses 3D CNN, dense connections, deep supervision together along with Dice objective function. Sub-acute stroke lesions have complicated features when compared to acute lesions [1]. To accurately segment both sub-acute and acute stroke lesions we propose Parallel Capsule Net that uses dense connections between parallel encoding pathways to minimize the information lost due to max pooling and enhance feature re-use. The proposed method uses a modified loss function which consists of a regional term (Generalized Dice loss + Focal Loss [8]) and a boundary term (Boundary loss) to address the problem of class imbalance.

## 2 Methodology

### 2.1 Proposed Architecture

In this paper, we propose a new encoder-decoder architecture (Fig. 1) which utilizes 3D convolutions, dense connectivity and parallel max pooling to efficiently train the model. The network has multiple encoding pathways (top to bottom) in parallel with dense connections across them (Fig. 3). We define an encoding block as a set of densely connected [7] convolutional layers whose outputs are concatenated as shown in Fig. 2. As we progress from the top to bottom, outputs from every convolutional layer in an encoding block are downsampled and passed on to the next encoding block. The outputs from the final encoding block are concatenated and given to the decoder. Instead of the traditional skip connections used in Unet [9], our architecture connects the encoder and the decoder through dense skip pathways [10]. In most of the encoder decoder architectures [9], downsampling is done by maxpooling. We introduce Parallel Max Pooling in our architecture to prevent loss of spatial information. Further details can be found in the Discussion. Table 1 summarizes information about the filters used in Parallel Capsule Net. After testing with different values for ‘number of filters’, we have found 20 filters for the first encoding block to be optimal (See Fig. 5).

**Table 1:** Details regarding the filters used in the architecture

| Block | Operation | Filter size | no. of filters |
| --- | --- | --- | --- |
| Encoder Block 1 | conv | 3x3x3 | 20 |
| Encoder Block 2 | conv | 3x3x3 | 40 |
| Encoder Block 3 | conv | 3x3x3 | 80 |
| Encoder Block 4 | conv | 3x3x3 | 160 |
| Decoder Block1 | 1 upsampling | 3x3x3 | 20 |
| Decoder Block2 | 2 upsampling | 3x3x3 | 40,20 |
| Decoder Block3 | 3 upsampling | 3x3x3 | 80,40,20 |
| Decoder Block4 | 4 upsampling | 3x3x3 | 160,80,40,20 |
| fully conv1 | conv | 1x1x1 | 10 |
| fully conv2 | conv | 1x1x1 | no. of classes |

**Fig. 1:**
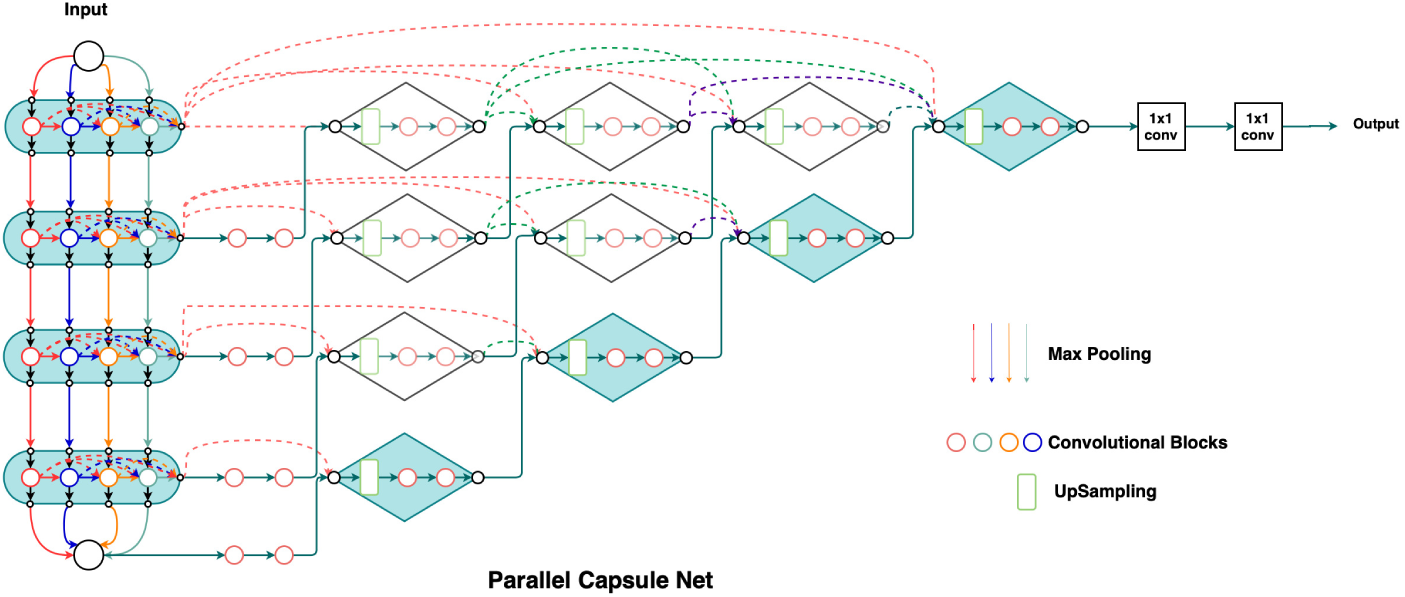
Parallel Capsule Net

**Fig. 2:**
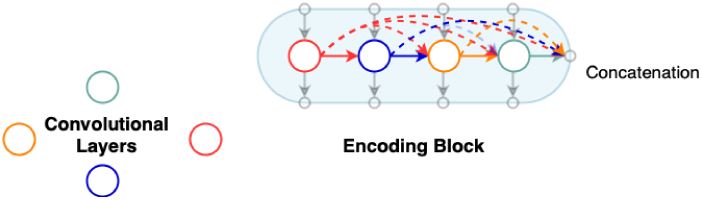
Dense Connections

**Fig. 3:**
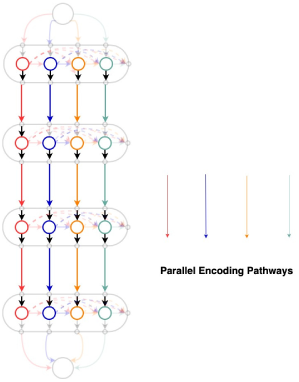
Parallel Pathways

### 2.2 Loss Function

Class imbalance in the field of medical image analysis is a common problem, where standard losses (cross entropy, Dice loss [12]) differ considerably across segmentation classes, which in turn affects training performance and stability. The use of Dice loss for unbalanced data would result in high precision, low recall segmentations which is undesired in computer aided diagnosis [13].

Focal loss (FL) [8], parametrized by γ, exponentiates the Dice loss(DL), helps to focus on hard classes which are detected with a lower probability. It reduces the contribution of background class and gives equal opportunity to the lesion class to learn efficiently. It is found that when γ *>*1, focal dice loss concentrates more on less accurate predictions which are misclassified.

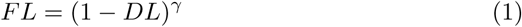

Boundary loss [15] concentrates on the distance metric which provides information complementary to the regional term and hence can mitigate the issue related to regional losses. Boundary loss uses the surface information of the ground truth via the level set function *ϕ*_*G*_ (*p*) (Fig. 4), which encodes the distance between each point p and *∂G*. The equation below summarizes the boundary loss where *s*_*θ*_(*p*) represents the softmax probability output of the network.

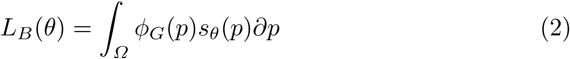

**Fig. 4:**
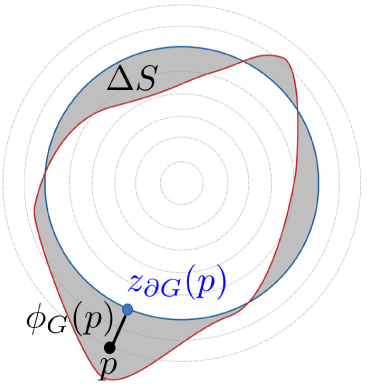
Boundary Integral

**Fig. 5:**
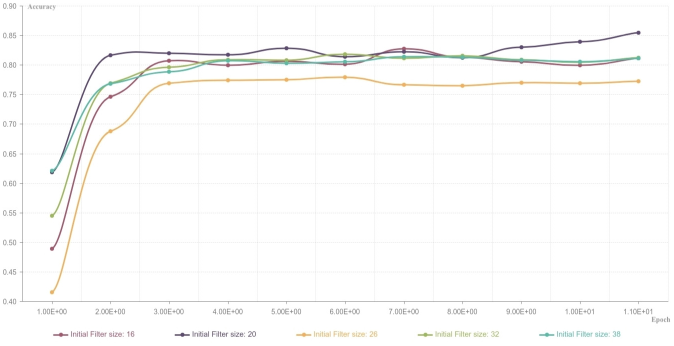
Comparison of different filters

We propose a modified loss function which consists of a regional term (dice loss and focal loss) and a boundary term (boundary loss) to leverage the advantages of the above mentioned losses. During the initial epochs, weights of the regional term are greater than the boundary term but as the training progresses, the weights of the regional term decrease polynomially while weights of the boundary term increase. The modified loss function is capable of handling class imbalance problem and balancing precision and recall effectively.

## 3 Experiment

### 3.1 Dataset

The architecture was trained on ISLES SISS 2015 [2], ISLES SPES 2015 [2] and ATLAS datasets [11]. ISLES SISS multi modal dataset contains 28 subjects for training and 36 subjects for testing. ISLES SPES multi modal dataset contains 30 subjects for training and 20 subjects for testing. ATLAS dataset contains T1 weighted images of 304 subjects. The manual ground truth labels were provided for the training samples. For the testing purpose, an expert radiologist was consulted to manually segment the test images. Detailed information about the dataset could be found on ISLES and ATLAS website [2] [11]. For preprocessing, N4 Bias field correction and [-1,1] normalization was used.

### 3.2 Training

During the training, 3D volumetric patches of size 32×32×32 with stride 8×8×8 were used. The use of patches reduces the memory requirement of the network and substantially increases the number of training samples, therefore removing the need for data augmentation by providing variance to the training input. Batch size of 8 was used for training. Adam Optimizer with a learning rate of 0.0001 was found to be best suited for training the proposed architecture. The network was trained for 20 epoch with a decay rate of 0.8 after every 5th epoch to improve the validation performance of the network. For the implementation purpose, TensorFlow was used and the experiments were run on a machine equipped with 4 NVIDIA GPUs with 16 GB of memory each.

## 4 Results

We compare in Table 2, the performance of our architecture with current state of the art algorithms, for three different datasets. The maximum scores achieved in each case are highlighted. As it can be seen, Parallel Capsule Net scores better than the other algorithms in all except one case.

**Table 2:** Comparison of Architectures on multiple Dataset.

| Dataset | Method | DSC(mean) | HD(mean) | Precision(mean) | Recall(mean) |
| --- | --- | --- | --- | --- | --- |
| ISLES 2015 SISS | DeepMedic [3] | 0.59 | 37.88 | 0.68 | 0.60 |
|  | Nested Unet [10] | 0.643 | 42.335 | 0.615 | 0.673 |
|  | Zhang et al. [1] | 0.58 | 38.98 | 0.60 | 0.68 |
|  | Anjali Gautam et al. [14] | 0.67 | 28.09 | 0.70 | 0.71 |
|  | **Parallel Capsule Net** | **0.754** | **22.871** | **0.739** | **0.769** |
| ISLES 2015 SPES<br>(Penumbra) | Nested Unet | 0.711 | 5.412 | 0.785 | 0.649 |
|  | McKinley et al. [2] | **0.82** | 1.65 | - | - |
|  | **Parallel Capsule Net** | **0.823** | **1.274** | **0.878** | **0.765** |
| ATLAS 2018 | Nested Unet | 0.89 | 12.122 | 0.924 | 0.857 |
|  | **Parallel Capsule Net** | **0.902** | **11.500** | **0.935** | **0.870** |

\* DSC: Dice Similarity Coefficient, HD: Hausdorff Distance

## 5 Discussion

It can be concluded from Fig. 6 that the algorithm works effectively for large and small lesions. Though our Dice Score is same as the current state of the art, the Hausdorff Distance is lower indicating accurate segmentation of boundaries of the penumbra that is important for clinical decision making. An example can be seen in Fig. 7. We have tested our model against stroke similar pathologies like small vessel ischemic disease and DWI artifacts. It can be seen from Fig. 8 that our model successfully detects the lesion and avoids segmenting stroke similar pathologies, which are a major challenge for other state of the art algorithms [2].

**Fig. 6:**
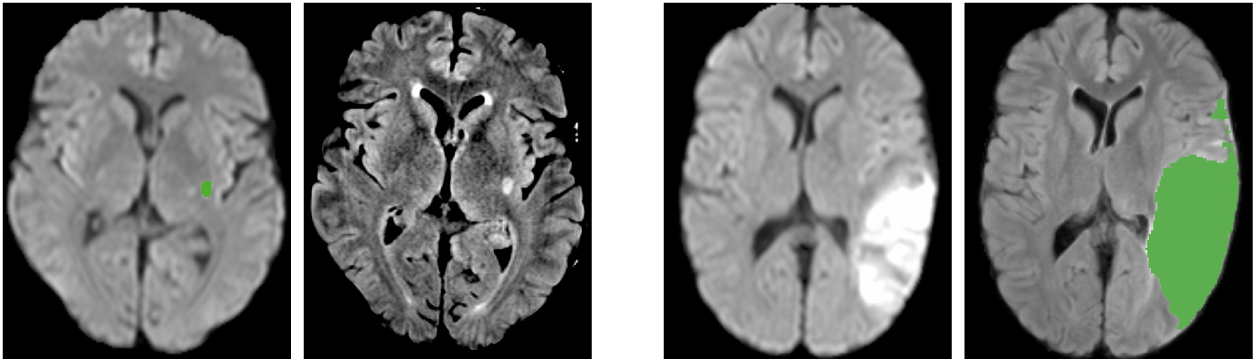
Case 1: DWI & Flair(with Segmented Label) in SISS.

**Fig. 7:**
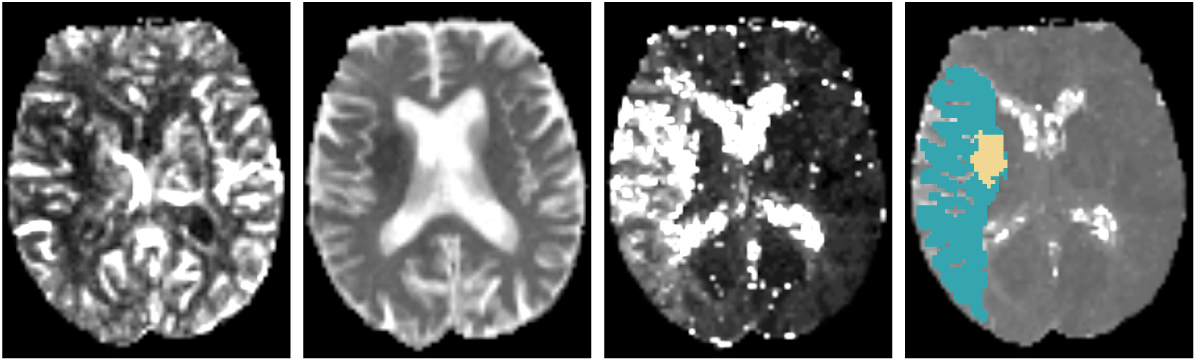
Case 2: CBF, DWI, TMax & TTP(with Segmented Label) in SPES

**Fig. 8:**
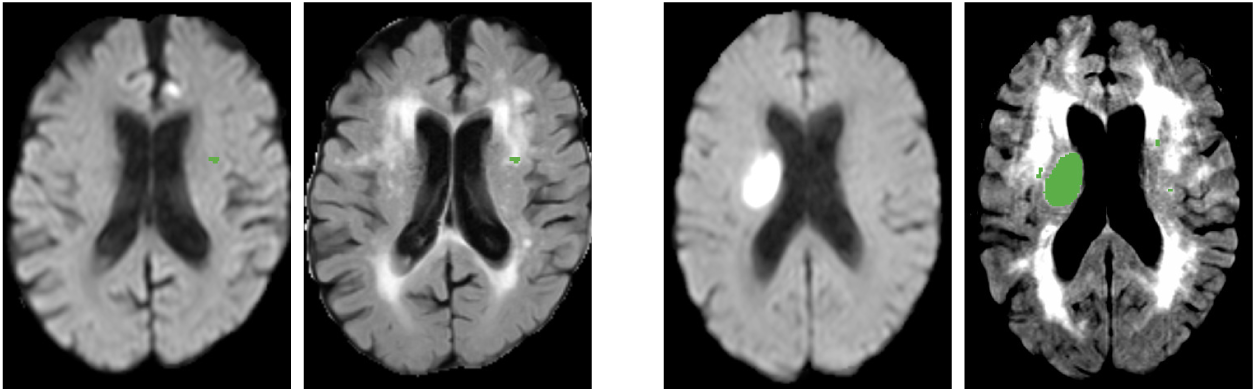
Case 3: Small Vessel Disease (DWI & Flair(with Segmented Label))

One of the novelty of the present algorithm is parallel maxpooling. In most of the encoder decoder architectures [9], downsampling is done by maxpooling to lower the memory footprint and enlarge the receptive field of the model. But, max pooling results in loss of spatial information, which might be detrimental in training. In Fig. 9&Fig. 10, an input is subjected to two convolutions and then downsampled. Each convolution feature map when maxpooled independently, retains a unique downsampled representation. By performing max pooling in every parallel pathway (Fig. 3), each pathway retains information, which may be lost in other pathways.

**Fig. 9:**
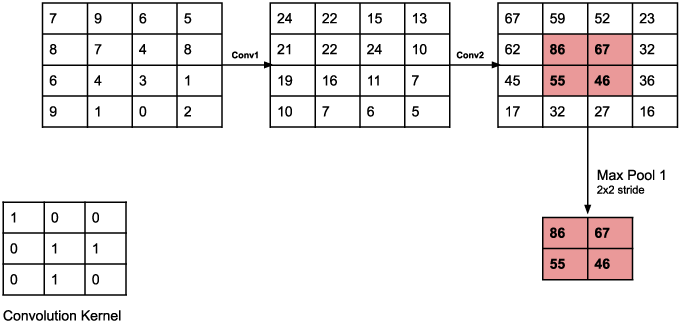
Single Max Pooling

**Fig. 10:**
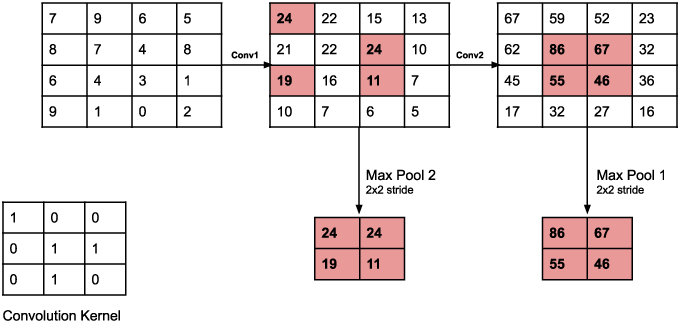
Parallel Max Pooling

Dense connectivity within an encoding block enhances feature re-use and promotes model compactness. Direct connections between layers improves the flow of information, causes a regularizing effect and prevents the problem of vanishing gradients [7]. Dense skip pathways reduce the semantic gap between the feature maps of encoder and decoder by introducing convolutional layers to skip connections. This enables the model to capture fine grained details of foreground.

Dense connections between the parallel pathways would combine various downsampled representations and facilitate collective flow of information. By introducing parallel pathways we also give network the flexibility to learn different feature representations independently. An important addition is the boundary term in the loss function. This is responsible for decrease in the Hausdorff distance in all the cases. Parallel Capsule Net seems to be promising in other medical segmentation tasks like tumor segmentation.

